# The realized host specificity of *Leptopilina japonica* and *Ganaspis kimorum*, adventive larval parasitoids of the invasive *Drosophila suzukii*

**DOI:** 10.64898/2025.12.22.696016

**Authors:** Paul K. Abram, Jessie Moon, Jessica L. Fraser, Jason Thiessen, Tara D. Gariepy

## Abstract

The non-native larval parasitoid wasps *Leptopilina japonica* and *Ganaspis kimorum* (Hymenoptera: Figitidae), which attack larvae of the invasive pest *Drosophila suzukii* (Matsumura) (Diptera: Drosophilidae), were unintentionally introduced to the Pacific Northwest of North America between 10 and 15 years ago, and are now well-established. Previous laboratory studies conducted to assess the ecological risk of classical biological control introductions suggested that *G. kimorum* is highly host-specific, whereas *L. japonica* can develop in several other drosophilid species. Information on realized host use of the parasitoids under field conditions in the context of host-parasitoid food webs in their non-native range is limited. A two-year field study in coastal British Columbia, Canada was conducted, using complementary sampling methods to document host-parasitoid associations and quantify parasitoid specificity with respect to drosophilid communities inhabiting ripe and rotting fruit. Consistent with previous laboratory findings, *G. kimorum* was reared exclusively from *D. suzukii*, while *L. japonica* emerged primarily from *D. suzukii* but also from *D. melanogaster* and, less frequently, from members of the *D. obscura* species group. *Leptopilina japonica* shared some host species with the resident parasitoid species *L. heterotoma* and *Asobara* cf. *rufescens*. A particularly striking result was the extent to which almost all drosophilid and parasitoid taxa in this study’s host-parasitoid trophic webs were non-native, complicating the interpretation of ‘non-target’ ecological effects of *L. japonica*. These results provide field-based evidence of realized host use by these two non-native larval parasitoids of *D. suzukii* that can inform their future use in classical and augmentative biological control programs.

## Introduction

The host specificity of parasitoids is a primary consideration when evaluating candidate agents for introduction in classical biological control programs against invasive insect pests. To minimize the potential for non-target impacts, parasitoids whose host range encompasses too many species other than the intended target for biological control are rejected during the process of their evaluation (van Driesche and Reardon 2004). Laboratory host range testing, where parasitoids are exposed to a range of potential “non-target” host species (i.e., those other than the target), are used to evaluate its fundamental host range; that is, the set of species the parasitoid will accept and which will support the development of the parasitoid’s offspring (van Lenteren et al. 2006). While these laboratory-based host range assessments are effective at identifying potential hazard, they tend to overestimate non-target host use and cannot predict population-level impact (Barratt et al. 1997; Haye et al. 2005; Cameron et al. 2013; van Driesche and Hoddle 2017; Haye et al. 2024). As a result, there is ongoing concern that some potentially effective biological control agents are being excluded from consideration based solely on overly cautious interpretations of lab-based host range data, without considering important ecological filters that may restrict parasitoid host range under field conditions (Heimpel et al. 2024; Abram et al. 2024).

At the same time, a growing number of unintentional introductions of biological control agents into regions outside of their native range are being discovered, sometimes even as they are undergoing formal evaluation for intentional release (Mason et al. 2017; Weber et al. 2021). These unintentional introductions offer an opportunity to observe the realized (i.e., ecological) host range of non-native natural enemies when exposed to new communities of hosts in natural settings (Haye et al. 2024). Comparisons of predicted versus actual non-target host use are still relatively rare for arthropod biological control agents (Barratt et al. 1997; Haye et al. 2005; Cameron et al. 2013; Haye et al. 2024), and additional case studies could help refine how host range data are interpreted during the evaluation of biological control agents. While relatively host-specific parasitoids introduced intentionally for classical biological control might be expected to form simple, modular food chains with their invasive hosts (Hawkins et al. 1999), unintentionally introduced parasitoid species with broader host ranges may integrate themselves more extensively into food webs (Hawkins and Marino 1997).

The larval parasitoids *Leptopilina japonica* Novkovic & Kimura (Hymenoptera: Figitidae) and *Ganaspis kimorum* Buffington (Hymenoptera: Figitidae; formerly ‘G1’ *G. brasiliensis* – see Sosa-Calvo et al. 2024; Hopper et al. 2024) were both evaluated as candidates for classical biological control of the globally invasive fruit pest *Drosophila suzukii* (Matsumura) (Diptera: Drosophilidae) (Stahl et al. 2024; Rossi Stacconi et al. 2025). Research on these parasitoids, conducted both in their native ranges (China, South Korea, and Japan) and in quarantine facilities in North America and Europe, began soon after *D. suzukii* invaded these continents in the late 2000s (Asplen et al. 2015; Daane et al. 2016; Girod et al. 2018a,b; Giorgini et al. 2019; Seehausen et al. 2020; Daane et al. 2021). Laboratory tests confirmed that the fundamental host ranges of both parasitoid species were restricted to the family Drosophilidae. However, while *G. kimorum* appeared to parasitize only *D. suzukii* and its close relative *D. melanogaster*, *L. japonica* was capable of attacking and developing in a broader set of drosophilid hosts. Based on these host range assessments, a release permit for *G. kimorum* was approved by regulators in the USA, Italy, and Switzerland (Stahl et al. 2024), whereas a permit for intentional release of *L. japonica* in the USA was denied (K. Daane, personal communication), and its introduction was not pursued elsewhere.

Nevertheless, beginning in 2019, both species were discovered to be present in the British Columbia (Canada) and Washington State (USA) (Abram et al. 2020; Beers et al. 2022), and were found to be well established (Abram et al. 2022a). Around the same time that *L. japonica* was first detected in Canada, it was also found in Italy, and has since been detected in other regions of North America and Europe (Gariepy et al. 2024; Rossi Stacconi et al. 2025). Meanwhile, *G. kimorum* has been released for classical biological control of *D. suzukii* in the USA, Italy, and Switzerland (Stahl et al. 2024) and is now being redistributed from western to eastern Canada (T.D. Gariepy, unpublished data). Given its narrow host range, *G. kimorum* is considered to pose minimal ecological risk (Stahl et al. 2024). In contrast, the broader fundamental host range of *L. japonica* has prompted calls for further evaluation of its realized host use in invaded areas, especially in the context of potential redistribution or augmentative use (Rossi Stacconi et al. 2025).

Recently, further research on *G. kimorum* and *L. japonica* has begun to clarify how laboratory host range predictions correspond to patterns of host use in the field. Field cage (Seehausen et al. 2022), olfactometry (Giorgini et al. 2024), and post-release field studies (Fellin et al. 2023) suggest that *G. kimorum* is indeed highly specialized on *D. suzukii* in nature. Preliminary field observations in Italy suggest that *L. japonica*, by contrast, may parasitize a wider range of drosophilid species (Fellin et al. 2023). However, how these two parasitoid species fit into food webs where they have been introduced is still not well understood – indeed, surprisingly little is known about drosophilid-parasitoid food webs in nature in many areas of the world (Lue et al. 2021; Bennett et al. 2024).

The objective of this study was to evaluate the realized specificity of *L. japonica* and *G. kimorum* under field conditions. Based on previous host range data, it was predicted that *G. kimorum* would be reared only from *D. suzukii*, while *L. japonica*, as an oligophagous parasitoid, would also be reared from other drosophilid groups but would show evidence of preference for *D. suzukii*.

## Materials and Methods

### Study sites and sampling design

The study was conducted over two years (2022 and 2023) in the eastern Fraser Valley of south coastal British Columbia, Canada, where *D. suzukii*, *L. japonica,* and *G. kimorum* are known to be common and widespread (Abram et al. 2020; Abram et al. 2022a; Capko et al. 2024). In 2022, parasitized Drosophilidae associated with rotting fruit in different habitats were collected at eight field sites. At two sites, < 10 parasitoids were reared from fruits, and so they were excluded from subsequent analysis. In 2023, sampling focused on a subset of three representative sites from which drosophilids and their parasitoids were consistently reared in 2022. To capture as much diversity as possible in drosophilid and parasitoid community composition, study sites included a variety of habitat types (Table 1). The parasitoids *L. japonica* and *G. kimorum* were already known to be present at several of the sites based on past studies by our research group (Abram et al. 2020; Abram et al. 2022a; Capko et al. 2024). In both years, naturally occurring rotting fruit and sentinel fruit bait traps was collected. In 2023, fresh ripening fruit was also collected, when possible. Standard laboratory incubation conditions for all the field collections documented below were 23 ± 1°C, 55-65% RH, and a 16:8h light:dark cycle.

**Table 1.**
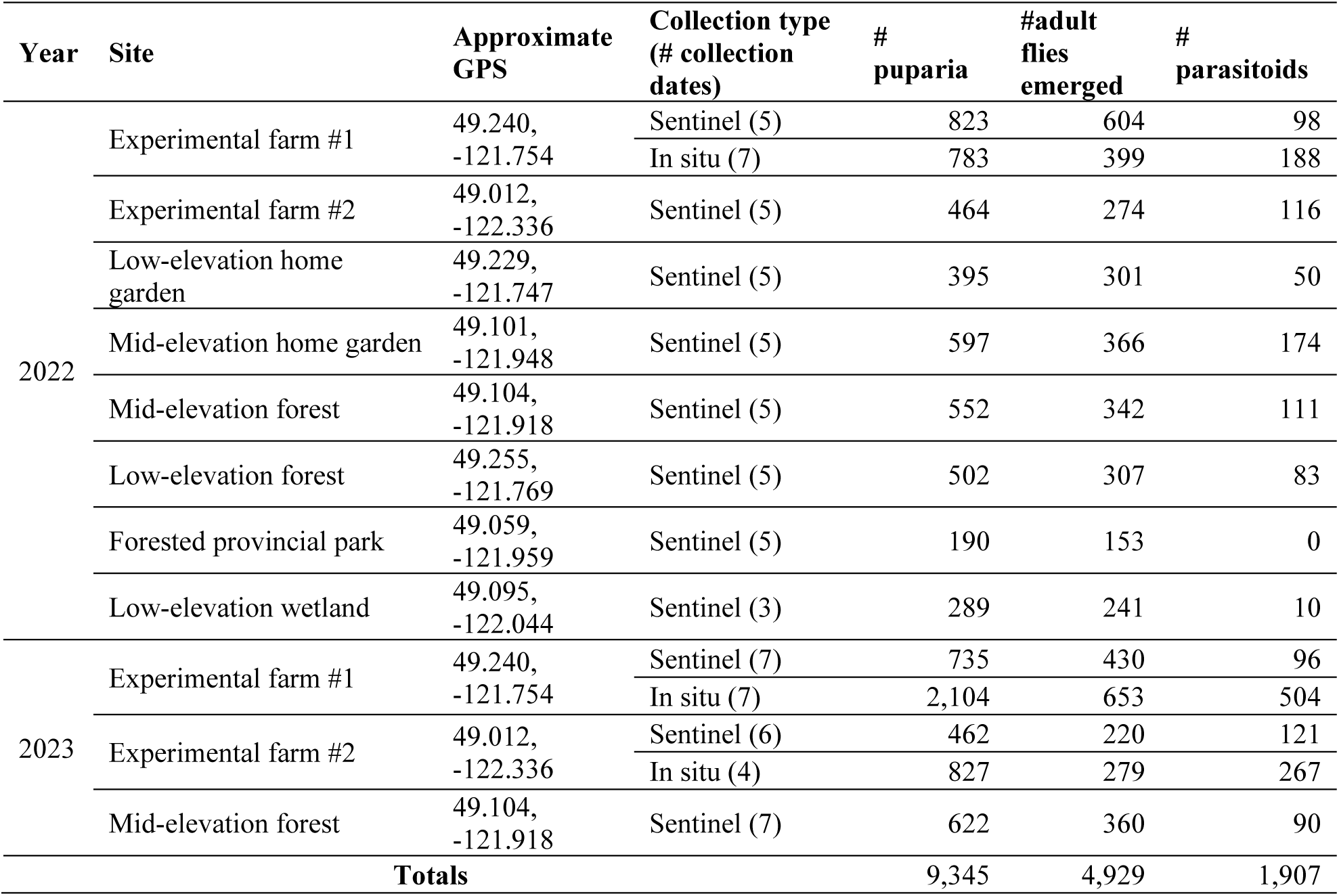
Study sites, collection types (sentinel fruit bait traps or *in situ* collections of rotting fruit from the ground) with the number of collection dates on which at least one drosophilid puparium was found, the total number of puparia reared (all collection dates pooled), and the number of adult drosophilid flies and parasitoids reared from the puparia (all collection dates pooled). Any puparia that did not result in the emergence of adult flies or parasitoids (i.e., # puparia – # adult flies emerged – # parasitoids) died during development.

### Collections of naturally occurring rotting fruit

To document *in situ* drosophilid larva-parasitoid associations, following Fellin et al. (2023), dropped and rotting (= on the ground, showing signs of decay) fruit were collected from the ground in the field. These fruits were expected to contain *D. suzukii* larvae (from prior oviposition into ripening fruit before it dropped) as well as larvae of other drosophilid species that subsequently lay their eggs in rotting fruit. In total, 18 individual collections of rotting fruit were made from two experimental farm sites over the two years (see Online Resource 1; Methods S1 for further details).

Dropped, rotting raspberry and blackberry fruits were collected from the ground by hand at intervals of 4-12 days, opportunistically, during different but overlapping seasonal periods each year (2022: 25 July to 19 September, 2023: 10 July to 16 August), up to a maximum of 200g of fruit per collection date per site. Fruits were returned to the laboratory, gently squashed in a plastic bag, and floated in 500mL of brown sugar solution so that drosophilid larvae would leave fruits and rise to the surface (Dreves et al. 2014) (Online Resource 1; Figure 6). *Drosophila* larvae of mid- to late-instar larval stages were then transferred with a fine paintbrush to ∼15 mL of artificial rearing medium (Drosophila 4-24 medium, Carolina, USA) in plastic vials (diameter: 2 cm; height: 9 cm) supplemented with 3-5 grains of baker’s yeast, closed with cellulose plugs with half of a Kimwipe (Kimtech, Canada) provided for the drosophilid larvae to pupate on. The medium used, especially when supplemented with yeast, is suitable for the development of a wide variety of *Drosophila* species, including *D. suzukii* and the other frugivorous species known to be present in the study area (e.g. Stratman et al. 1998; Larsson et al. 2004; Kokate et al. 2022; Nixon et al. 2022; P. Abram, personal observations). A maximum of 100 larvae were transferred to each vial. Some samples contained thousands of larvae and it would not have been practical to rear them all; in these cases a subset of up to 500 was taken. Two times weekly, vials were checked for the presence of drosophilid puparia.

In the first year of the study (2022), parasitoid diversity and host-parasitoid associations were documented, but not per-drosophilid-species parasitism levels Thus, the puparia retrieved from the diet tubes were incubated in groups of up to 50 in Petri dishes (diameter: 5 cm; height: 0.9 cm) lined with filter paper moistened with water until unparasitized puparia emerged as flies, which were removed and preserved in 95% EtOH. The remaining unemerged puparia were considered potentially parasitized. These were individually photographed to document puparial morphology, and then were incubated individually in 1.5 mL microcentrifuge tubes containing moistened (with water) cotton until parasitoid emergence (or up to 40 days if no parasitoid emerged). Puparia containing dead, unemerged parasitoids were dissected and the parasitoid identified, if possible. If identification was not possible, the parasitoid was not included in the analysis of parasitoid-host associations but was included in calculations of parasitism levels. Drosophilid puparia were identified to species or species group levels, following the methods of Abram et al. (2022b), with the identification of a subset of them (n= 399) confirmed with DNA barcoding (Online Resource 1; Methods S1). Identifications based on puparial morphology were 98-100% accurate (Online Resource 1; Methods S1). All barcoded *D. melanogaster* group puparia were *Drosophila melanogaster* Meigen, whereas *D. obscura* group puparia were 89% *Drosophila subobscura* Collin and 11% *Drosophila affinis* Sturtevant. Parasitoids were identified following the key of Abram et al. (2022b), with a subset of the representative voucher specimens confirmed by Dr. Matt Buffington and Dr. Robert Kula (United States Department of Agriculture, Washington, DC). Voucher specimens have been retained at the Agassiz Research and Development Centre, Agassiz, British Columbia, Canada and deposited in the Canadian National Collection of Arthropods and Nematodes.

In 2023, in addition to documenting drosophilid-parasitoid associations, percentage parasitism of each drosophilid species’ (or species group’s) puparia collected on each date was determined. Here, the puparia retrieved from the diet tubes were sorted by morphology before being incubated in Petri dishes. This allowed the number of unparasitized drosophilids and parasitoids emerging from each type of puparium to be documented for each collection. Unemerged parasitoids were dissected out of puparia and identified when possible.

### Sentinel fruit bait traps

The spatial and temporal scope of the collections of naturally occurring rotting fruit was relatively limited in space (number of sites) and time (during the season), and overwhelmingly contained *D. suzukii* larvae rather than the other species of drosophilids. Thus, an additional and complementary method to determine drosophilid larva-parasitoid associations was used that, while less realistic than collections of naturally occurring fruit, overcame these limitations. This involved deploying sentinel fruit baits at regular intervals during each of the two study years.

Bait traps were designed to initially contain fresh fruit that would be attractive to *D. suzukii* and its larval parasitoids, and subsequently (once the fruit naturally rotted) other drosophilids and their associated larval parasitoids. The traps also needed to have openings large enough so that the entry of drosophilids and their parasitoids was not obstructed and some ventilation was permitted, but strong enough to protect the fruit from vertebrate wildlife. The traps (Online Resource 1; Figure 6) consisted of red metal wire mesh birdfeeders (14.5 × 14.5 × 14.5 cm; mesh size: 1 × 0.5 cm; NO/NO feeders, Amazon, Canada) containing fruit purchased at local grocery stores. In each trap, 25 blackberries and two bananas were placed, and paper towel was placed in the bottom of the trap to absorb excess liquid and prevent the fruit from dropping through the holes on the underside of the trap. Blackberries were chosen based on their known attractiveness to *D. suzukii* and its parasitoids in the study region (e.g., Abram et al. 2022a), whereas bananas are a popular substrate for trapping a variety of other drosophilid species (e.g., Da Cunha et al. 1951; Daane et al. 2016). Traps were hung on vegetation, as close as possible to naturally occurring fruiting plants, at study sites between 1 and 2 meters above the ground. Shaded locations were chosen to prevent premature fruit desiccation as much as possible. Traps were set out for long enough to allow oviposition and larval development of drosophilid larvae but before larvae pupated; between 100 and 120 degree-days (base 7.2°C), with local weather data (Environment and Climate Change Canada 2023) (Tochen et al. 2014). New traps were set up and taken down (always in the same locations) between 6 and 7 times per site per year during the seasonal activity period of *L. japonica* and *G. kimorum* in the study region, between May and October (2022: May 11–October 4; 2023: May 31–August 9). One trap per site per date was set out in the first year of the study, and three traps per site (placed 50–100m apart) were set out in the second year of the study to potentially capture more within-site variation in drosophilid and parasitoid communities. Fruit was returned to the laboratory and drosophilid larvae were extracted from samples and reared to determine host-parasitoid associations in the same way as described above for naturally occurring rotting fruit in each study year. The number of successful samples per site varied because on some collection dates and some sites, no drosophilid larvae were extracted after collection.

In the second year of the study, to confirm that *G. kimorum* was present at the study sites during the period where sentinel bait traps were deployed, opportunistic collections were made of between 230 and 822 g (mean ± SE: 538 ± 33 g; n = 30) of ripe but undamaged fruit from fruiting plants between 8 and 9 times at each of the three study sites (Online Resource 1; Table 3). This fruit was incubated and parasitoids were reared from the fruit in plastic, ventilated containers under standard rearing conditions (Abram et al. 2022a) to document whether *G. kimorum* was present at each site.

### Food web construction and quantitative analysis of parasitoid specificity

For both years of the study, rearing data was used to represent host-parasitoid associations in quantitative food webs, considering parasitized hosts only (Sunderland et al. 2023), for each unique combination of site, year, and collection method (dropped rotting fruit or sentinel bait traps).

We quantified realized host specificity for each parasitoid species using the Paired Difference Index (PDI), a metric of food web specialization, following Ramirez et al. (2022). PDI evaluates the disparity between the strongest host interaction and all remaining interactions, yielding a measure of preference structure within quantitative food webs (Poisot et al. 2012). PDI values range between 0 (absolute generalist) to 1 (absolute specialist). Analyses were restricted to parasitoid-host interactions observed in 2023 (when the abundance of unparasitized puparia of each host was measured; see above), using data from *in situ* rotting fruit collections and bait traps. Observed values of PDI were compared with expectations generated from a null model (10,000 iterations) built using *econullnetr* (Vaughan et al. 2018). The null model weighted interaction probabilities by their relative abundances. These simulations produced sampling distributions from which expected values, 95% confidence intervals, and standardized effect sizes (SES) were calculated. Host availability for each parasitoid species was quantified at the site-date-method level, such that the relative abundance of hosts in a given null model iteration reflected the composition of hosts actually present at the same site sampled with a given method on the same sampling date. This ensured that null expectations were conditioned on the pool of hosts available to foraging parasitoids. Preference plots were also created that compare observed interaction frequencies for each host against the 95% confidence intervals from the null model (Vaughan et al. 2018; Ramirez et al. 2022). The food web metric calculations were implemented using the R package *bipartite* (Dormann et al. 2008), and all analyses were conducted in R (R Core Team 2024).

## Results

### Collections of naturally occurring rotting fruit

A total of 19 collections of rotting raspberry and blackberry fruit were done over the two years of the study (Table 1). A total of 3,714 drosophilid puparia extracted from these samples, of which 25.8% were parasitized. In both years, the vast majority of the parasitized puparia extracted from rotting fruits were *D. suzukii* (2022: 91.5%, 2023: 97.6%). Parasitized puparia of *Drosophila melanogaster* species group (2022: 5.8%, 2023: 2.3%) and *D. obscura* species group (2022: 2.7%, 2023: 0%) were much less frequently recovered from these samples.

Four species of parasitoids were reared from these puparia in both years: *L. japonica*, *Asobara* cf. *rufescens* (Förster) (Hymenoptera: Braconidae), *Leptopilina heterotoma* (Thomson) (Hymenoptera: Figitidae), and *G. kimorum* (Figure 1). *Leptopilina japonica* accounted for a large majority (2022: 86.8%, 2023: 90.1%) of the parasitoids reared from *D. suzukii* and was also reared from *D. melanogaster* group puparia (2022: 18.1%, 2023: 72.2% of all parasitoids of this species group) and *D. obscura* group puparia (2022: 20.0%). *G. kimorum* only emerged from *D. suzukii* puparia, albeit only accounting for small percentages (2022: 1.6%, 2023: 6.1%) of all parasitoids reared from this host species. *Asobara* cf. *rufescens* was reared from *D. suzukii*, *D. obscura* group, and *D. melanogaster* group puparia (Figure 1). *Leptopilina heterotoma* was only reared from *D. melanogaster* group and *D. obscura* group puparia (Figure 1).

**Figure 1.**
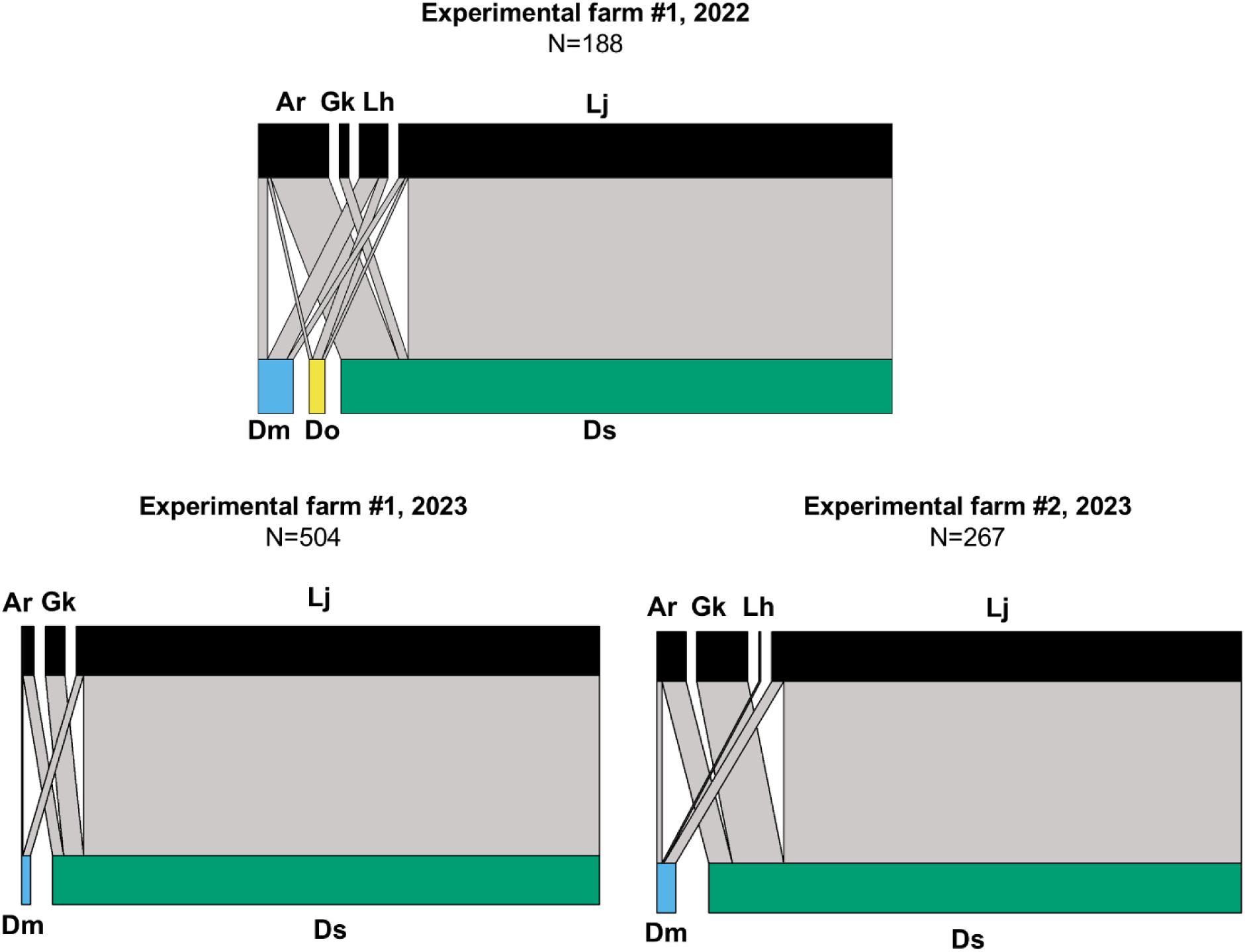
Quantitative food webs (parasitized puparia only; sample sizes, N, shown in titles) showing which species of parasitoids were reared from the puparia of different drosophilid species or species groups collected from rotting fruits in the two years of the study. The width of each bar shows the relative number of each parasitoid (upper bars on each food web) or host (lower bars). The width of the grey connecting lines, which are drawn between the parasitoid species and the hosts they were reared from, shows how many individuals of each parasitoid was reared from each species or species group of drosophilid. Species abbreviations: Ar – *Asobara* cf. *rufescens*; Gk – *Ganaspis kimorum*; Lj – *Leptopilina japonica*; Lh – *Leptopilina heterotoma*; Dm – *D. melanogaster* species group other than *D. suzukii*; Do – *D. obscura* species group; Ds – *Drosophila suzukii*.

In the second year of the study when percent parasitism levels were measured, 26.8% of *D. suzukii* puparia in rotting fruit were parasitized, with parasitism levels fluctuating between 4.2% and 44.2% among sites and collection dates (Online Resource 1; Figure 7). Of all the *D. melanogaster* species group puparia, 14.6% (8.0 –100.0%) of puparia were parasitized, whereas none of the *D. immigrans* puparia were parasitized (Online Resource 1; Figure 7).

### Sentinel bait traps

A total of 5,562 drosophilid puparia were extracted from fruit bait sentinel traps over the two years of the study. Identifiable parasitoids were reared or dissected from a total of 17.1% of these puparia (Table 1). Parasitized puparia belonged to, in decreasing order of prevalence: the *D. melanogaster* species group (2022: 49.5%, 2023: 46.6%), the *D. obscura* species group (2022: 41.7%, 2023: 36.5%), *D. suzukii* (2022: 3.9%, 2023: 13.7%), and *D. immigrans* (2022: 4.8%; 2023: 3.3%).

*Leptopilina heterotoma* was the most common parasitoid species in these collections in the first year of the study (73.7% of all parasitoids), whereas *L. japonica* was the most common parasitoid in the second year (54.4%) (Figures 2, 3). *Asobara* cf. *rufescens* made up a small percentage of all parasitoids in both years (2022: 1.1%, 2023: 1.0%). *Ganaspis kimorum* was entirely absent from these samples.

**Figure 2.**
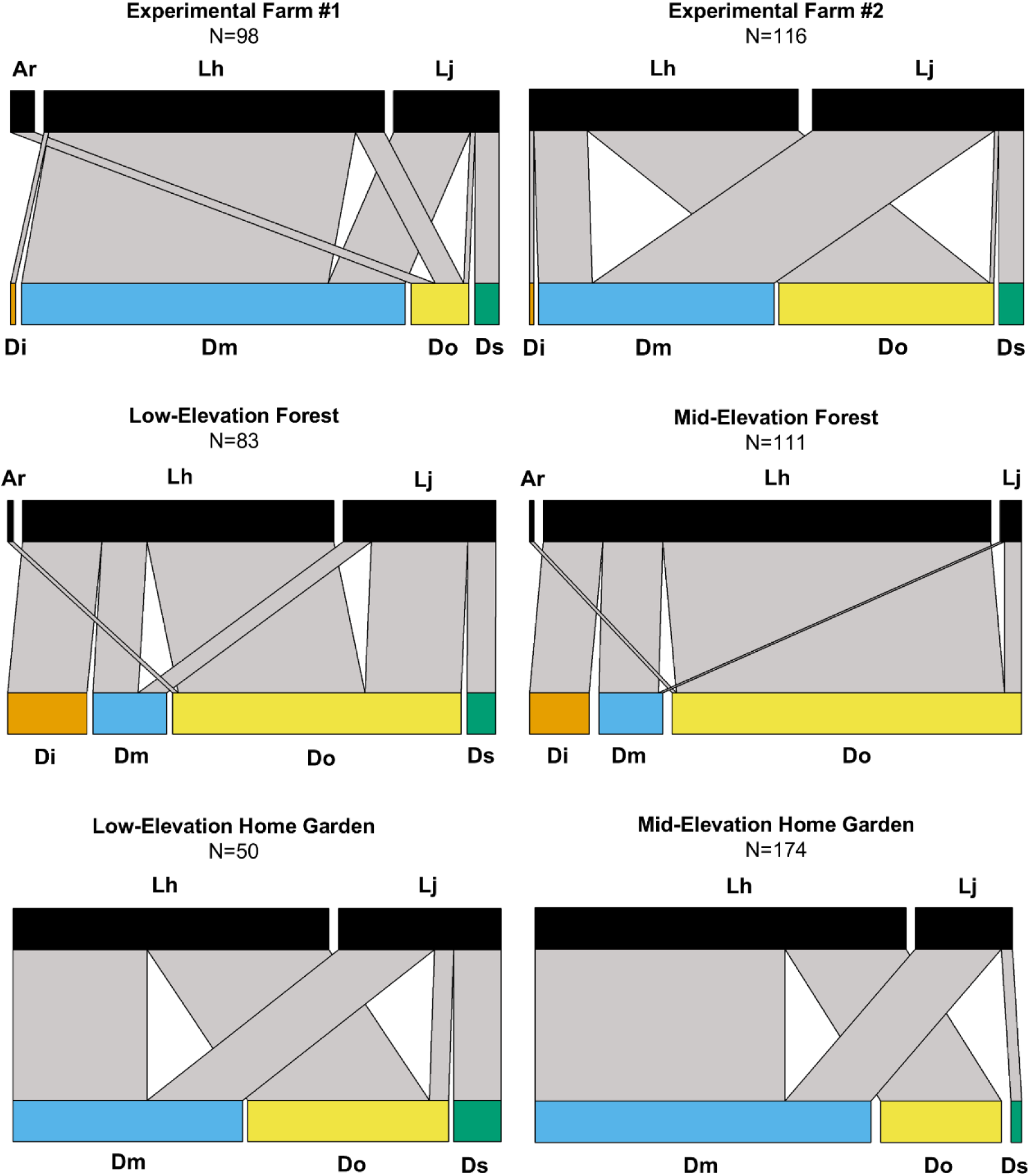
Quantitative food webs (parasitized puparia only; sample sizes, N, shown in titles) showing which species of parasitoids were reared from the puparia of different drosophilid species or species groups collected from sentinel fruit bait traps 2022, the first year of the study. The width of each bar shows the relative number of each parasitoid (upper bars on each food web) or host (lower bars). The width of the grey connecting lines, which are drawn between the parasitoid species and the hosts they were reared from, shows how many individuals of each parasitoid was reared from each species or species group of drosophilid. Total sample sizes are given in Table 1. Species abbreviations: Ar – *Asobara* cf. *rufescens*; Lj – *Leptopilina japonica*; Lh – *Leptopilina heterotoma*; Dm – *D. melanogaster* species group other than *D. suzukii*; Do – *D. obscura* species group; Ds – *Drosophila suzukii,* Di – *Drosophila immigrans*.

**Figure 3.**
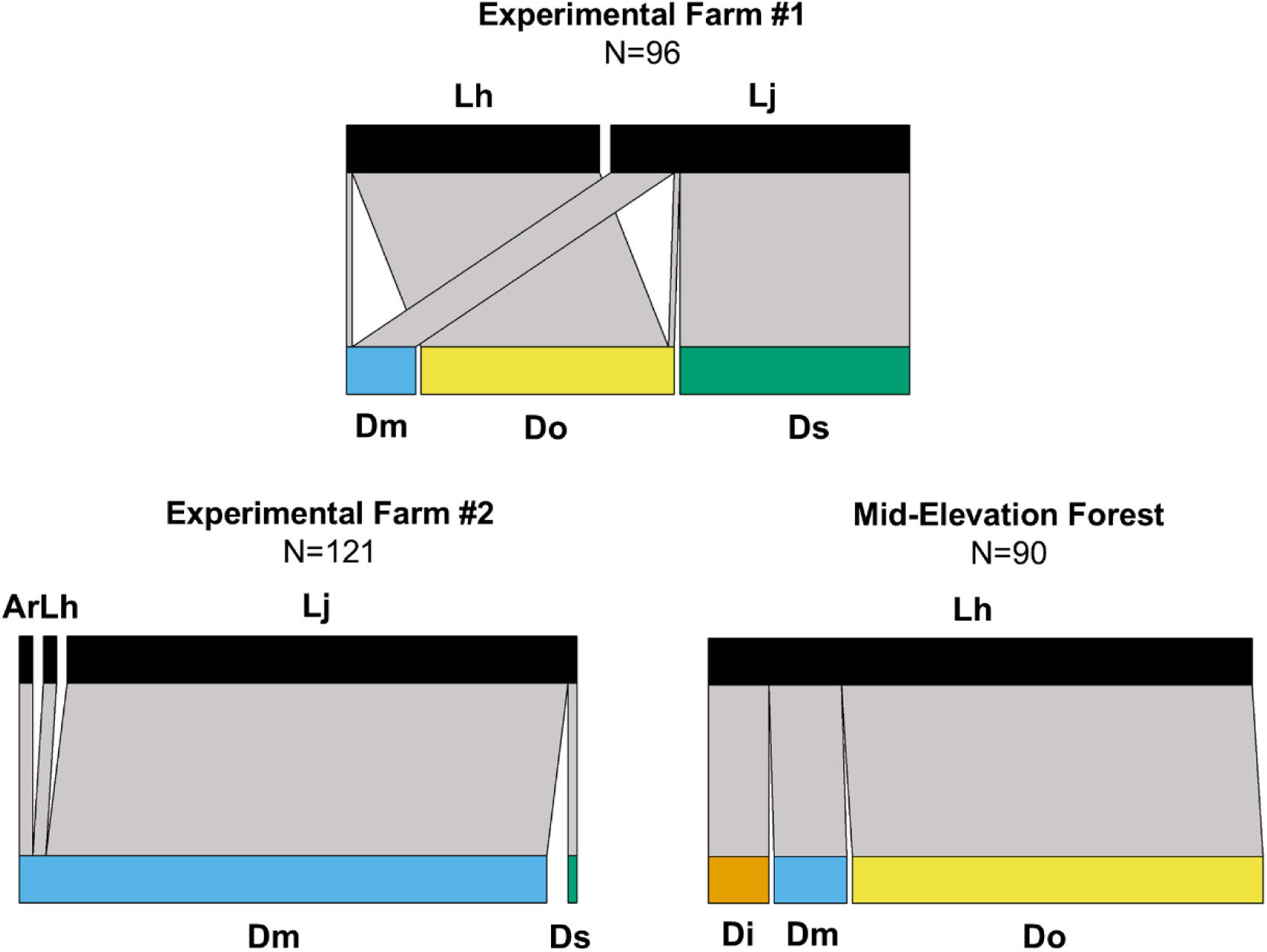
Quantitative food webs (parasitized puparia only; sample sizes, N, shown in titles) showing which species of parasitoids were reared from the puparia of different drosophilid species or species groups collected from rotting fruit in the second year of the study (2023). The width of each bar shows the relative number of each parasitoid (upper bars on each food web) or host (lower bars). The width of the grey connecting lines, which are drawn between the parasitoid species and the hosts they were reared from, shows how many individuals of each parasitoid was reared from each species or species group of drosophilid. Total sample sizes are given in Table 1. Species abbreviations: Ar – *Asobara* cf. *rufescens*; Lj – *Leptopilina japonica*; Lh – *Leptopilina heterotoma*; Dm – *D. melanogaster* species group other than *D. suzukii*; Do – *D. obscura* species group; Ds – *Drosophila suzukii,* Di – *Drosophila immigrans*.

*Leptopilina japonica* was present in 8/9 site-years and was the only parasitoid reared from *D. suzukii* puparia (Figures 2, 3). *Leptopilina japonica* was also reared from the *D. melanogaster* species group (0.0–94.9% of all its parasitoids; *x̄* = 44.0%; 8/9 site-years where this host was present) and *D. obscura* species group (0.0–9.5%; *x̄* = 7.5%; 6/9 site-years). *Leptopilina heterotoma* was overall the dominant parasitoid of the *D. melanogaster* species group (2.5-100.0%, *x̄* = 55.7%, 9/9 site-years), the *D. obscura* species group (50.0–100.0%, *x̄* = 86.9%, 8/8 site-years), and *D. immigrans* (100%, 5/7 site-years) (Figures 2, 3). *Asobara* cf. *rufescens* was present in 4/9 site-years and made up a minority (0.0–41.6%) of the parasitoid complex of the *D. melanogaster* and *D. obscura* species groups from bait traps.

In the second year of the study when percent parasitism levels were measured, 11.6% of all *D. suzukii* puparia in bait traps were parasitized, with parasitism levels fluctuating between 0 and 45% among sites and collection dates (Online Resource 1; Figure 8). Overall, 47.4% (0.0–100.0%) of *D. melanogaster* species group puparia, 28.2% (0.0–47.4%) of the *D. obscura* species group, and 1.69% (0.0–4.5%) of *D. immigrans* puparia were parasitized.

In the second year of the study, *G. kimorum* was reared from naturally occurring ripening fruit at all three study sites where sentinel fruit bait traps were deployed (Online Resource 1; Table 3), confirming its presence at these sites despite its absence in sentinel fruit bait traps. *Leptopilina japonica* was also present in ripening fruit samples at all three sites.

### Quantitative analysis of parasitoid specificity

Specialization (as indicated by PDI values) for *L. japonica* and *L. heterotoma* were significantly higher than expected under the null model, indicating greater specialization (fewer host species attacked) than predicted by random host use (Table 2). In contrast, the PDI values for *G. kimorum* and *A.* cf. *rufescens* did not differ from the null expectation, indicating that their observed host breadth was consistent with expectations based on host availability and sampling effort. *Leptopilina japonica* was reared more often from *D. suzukii* and less often from *D. immigrans* than the null model predicted (Figure 4); in contrast, its parasitism of *D. obscura* species group and *D. melanogaster* were consistent with frequency-dependent host use. *Leptopilina heterotoma*, in contrast, was reared less often from *D. suzukii* and *D. immigrans* than expected from the null model, and reared more often than under null expectations from *D. obscura* species group and *D. melanogaster* (Figure 4). The host species use of *A.* cf. *rufescens* and *G. kimorum* was indistinguishable from frequency-dependent use of hosts within the samples from which they were reared.

**Figure 4.**
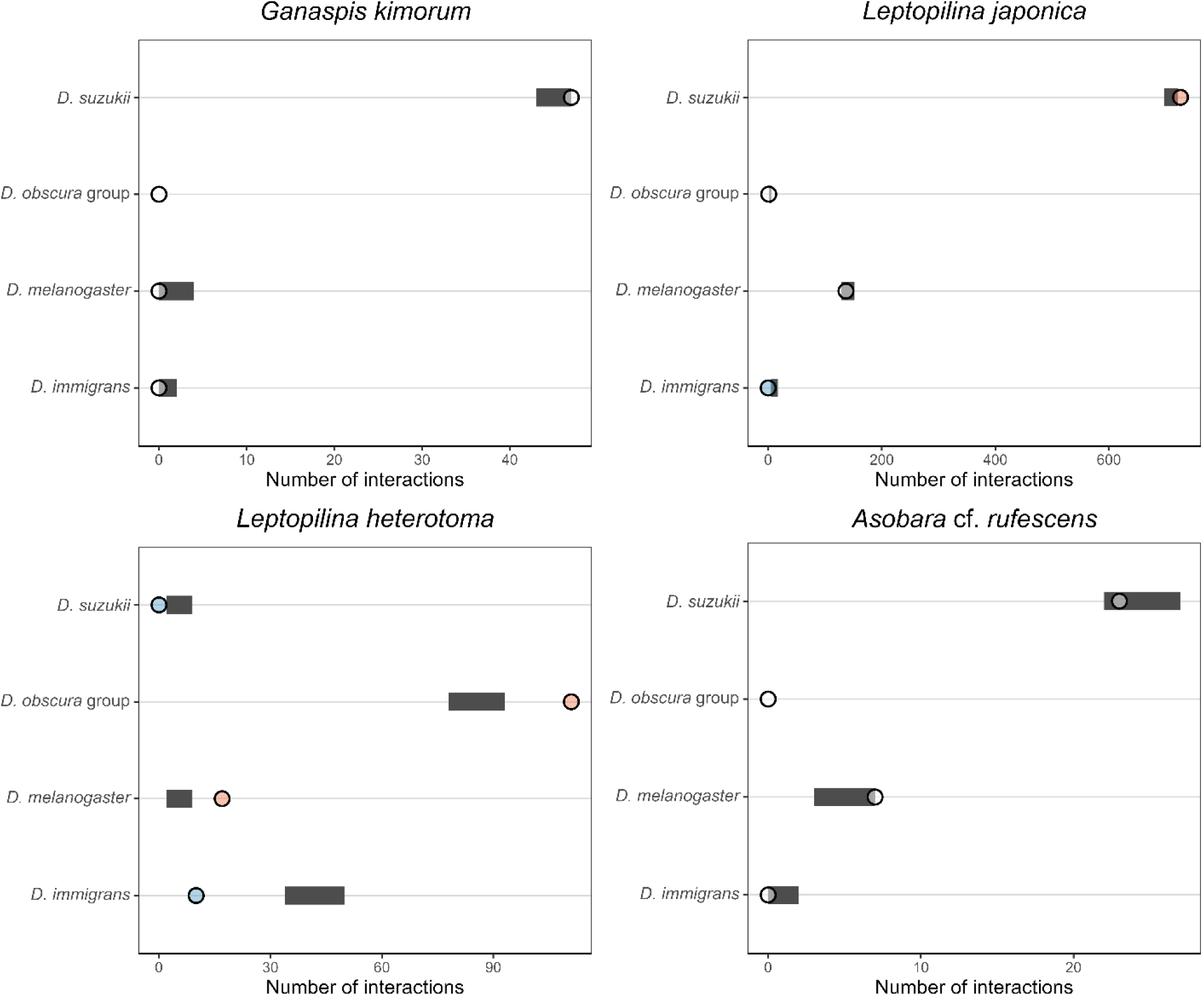
Preference plots for four parasitoid species reared from *Drosophila* hosts in 2023 bait trap and *in situ* rotting fruit field collections. For each parasitoid, panels show the observed frequency of emergence from each host species (points) compared with the 95% range of interaction strengths predicted by a null model that accounts for host availability and sampling effort (horizontal grey bars). Orange points to the right of the null range indicate host use stronger than expected, blue points to the left indicate weaker-than-expected interactions, and white points falling within the interval indicate no detectable deviation from random host use.

**Table 2.** Paired Difference Index (PDI) for four larval parasitoid species attacking *Drosophila* hosts from data collected from *in situ* rotting fruits and bait traps in 2023. Observed values for PDI (0 – absolute generalist, 1 – absolute specialist) are compared with expectations generated from a null model, along with their 95% confidence limits (CL) and standardized effect sizes (SES). The ‘Test’ column indicates whether the observed specialization is significantly higher, lower, or not significantly different (ns) from that expected under the null model.

| Species | Observed PDI | Expected PDI | Lower CL | Upper CL | Test | SES |
| --- | --- | --- | --- | --- | --- | --- |
| *G. kimorum* | 1.00 | 0.99 | 0.97 | 1.00 | ns | 1.39 |
| *L. japonica* | 0.94 | 0.93 | 0.92 | 0.93 | Higher | 2.74 |
| *L. heterotoma* | 0.92 | 0.79 | 0.75 | 0.84 | Higher | 5.40 |
| *A. cf. rufescens* | 0.90 | 0.93 | 0.88 | 0.96 | ns | -1.37 |

## Discussion

The results largely validated what was predicted by prior laboratory and semi-field studies. *Ganaspis kimorum* only parasitized *D. suzukii*, whereas *L. japonica* does parasitize other species of drosophilids in nature, although it demonstrates a tendency to parasitize *D. suzukii* over other available host species. A particularly striking result was that the drosophilid-parasitoid food webs sampled almost entirely consisted of non-native species at both trophic levels (summarized in Figure 5). The implications of the findings in terms of parasitism of *D. suzukii*, parasitism of other drosophilid species, and food web interactions among parasitoids sharing hosts are discussed below.

**Figure 5.**
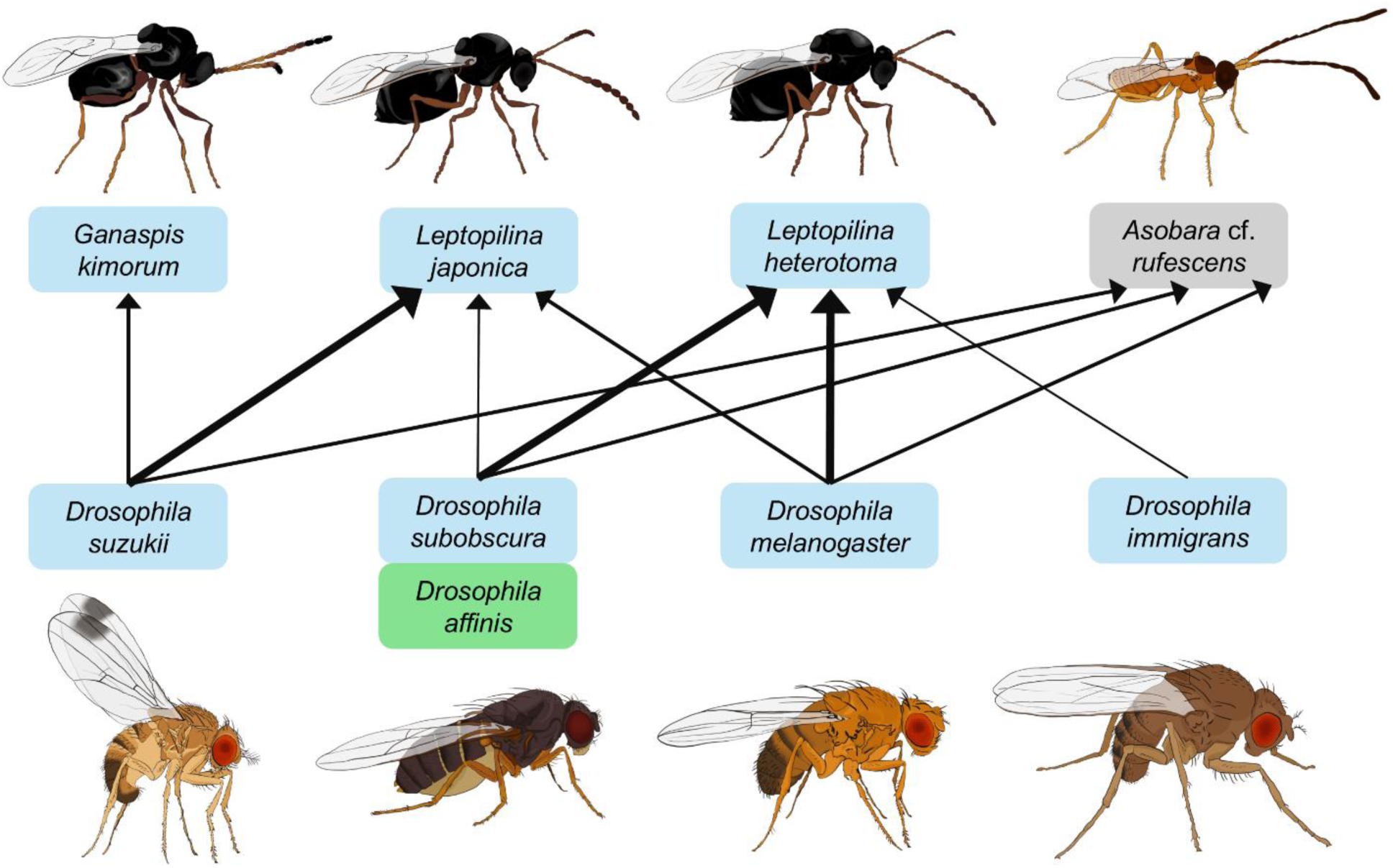
A generalized food web of drosophilid-larval parasitoid associations within ripening and rotting fruit in south coastal British Columbia, Canada based on the results of this study. The arrows show the direction of energy flow. The relative thickness of the lines and arrows shows, as per the analysis in Figure 4, relative preference of parasitoids for different host species: i.e., whether parasitism of a host species by a parasitoid species was less-than-frequency-dependent (thinnest arrows), frequency-dependent (medium arrows), or more-than-frequency dependent (thickest arrows). Species in blue boxes represent non-native or cosmopolitan species; green represents native species; grey represents species of unknown biogeographic origin. The two species in the *D. obscura* species group (*D. subobscura* and *D. affinis*) are shown with a single link to each parasitoid as the methodology did not allow per-species parasitoid associations to be determined within this species group.

### Parasitism of D. suzukii

Past studies found that in areas where *L. japonica* is established, it tends to be the dominant parasitoid of *D. suzukii* in fresh (Abram et al. 2020; Abram et al. 2022a; Beers et al. 2022; Gariepy et al. 2024; Rossi Stacconi et al. 2025) and rotting (Fellin et al. 2023) fruit. In the present study region, *L. japonica* and *G. kimorum* are adventive and well established across a variety of different habitats and host plants in ripening fruit containing *D. suzukii* (Abram et al. 2020; Abram et al. 2022a; Beers et al. 2022). However, it was not known to what degree either species would be associated with rotting fruit. In the current study, *L. japonica* was usually the dominant parasitoid of *D. suzukii* in both *in situ* rotting fruit and sentinel fruit baits. *Ganaspis kimorum* was only reared from *D. suzukii* larvae inhabiting *in situ* rotting fruit, despite its presence at most study sites and that *D. melanogaster* larvae (which *G. kimorum* uncommonly parasitized in laboratory studies) were abundant. Given that *G. kimorum* is attracted to volatiles of ripening (but not rotten) fruit infested by *D. suzukii* (Giorgini et al. 2024), the presence of *G. kimorum* in *D. suzukii* larvae *in situ* rotting fruit likely resulted from attacks of early-instar *D. suzukii* larvae in ripening fruit before they fell to the ground and rotted. This also may be true of at least some of the *L. japonica* reared from the *in situ* rotting fruit. However, in the sentinel bait traps, which likely begin to decay rapidly after deployment, *G. kimorum* was not ever reared from *D. suzukii* larvae, whereas *L. japonica* was consistently present parasitizing *D. suzukii* in these samples. This finding, combined with the fact that *L. japonica* was reared from other drosophilid species that colonize fruit after it begins to rot (see below), suggests that *L. japonica* may forage on both ripening and rotting fruit in nature. That *L. japonica* forages for hosts on rotting fruit suggests that additional biological control of *D. suzukii* may occur in rotting fruit and that sampling ripening fruit may underestimate parasitism. However, it also creates an opportunity for *L. japonica* to parasitize other drosophilid species which feed on rotting fruits.

### Parasitism of other drosophilid species

Laboratory and semi-field studies done during the pre-release risk assessment of *L. japonica* and *G. kimorum* (Girod et al. 2018b; Daane et al. 2021; Seehausen et al. 2020, 2022) predicted that *G. kimorum*’s host range would be limited to *D. suzukii* in the field, whereas *L. japonica* may attack other drosophilid species, particularly in the *D. melanogaster* and *D. obscura* species groups. The results here show that these predictions were accurate, at least as applied to this study’s regional and ecological context. *Ganaspis kimorum* was never reared from any species of drosophilid other than *D. suzukii*, whereas *L. japonica* parasitized the two groups of non-target species that were found to be attacked in previous laboratory studies. However, the level of parasitism differed markedly between the two groups of drosophilids. *Leptopilina japonica* was commonly reared from *D. melanogaster* across sampling techniques, study sites and years. In fact, in three out of nine site-years, *L. japonica* was the dominant parasitoid of *D. melanogaster* in sentinel fruit bait traps. In contrast, while *L. japonica* was reared from *D. obscura* species group larvae, it was relatively uncommon: *L. japonica* was reared from *D. obscura* group larvae in a minority of collections, and parasitized less than 0.25% of *D. obscura* species group larvae in bait traps in the second year of the study when percent parasitism was measured. While caution should be exercised when relating point measurements of percent parasitism to true biological control impact (Van Driesche 1983), and the types of sentinel fruit baits used may not realistically represent what takes place in naturally occurring substrates, these results provide a preliminary indication that parasitism of *D. melanogaster* by *L. japonica* may potentially affect its abundance, while the population-level impact of *L. japonica* parasitism on *D. obscura* group drosophilids is likely to be minimal.

Analyses of specialization metrics provided additional insight into how sampling context shaped apparent host use. Both *L. japonica* and *L. heterotoma* exhibited significantly higher specificity (PDI values) than expected under the null model, reflecting their tendency to parasitize a subset of the host species available at a given site-date. More specifically, *L. japonica* was more likely to parasitize *D. suzukii* over other available hosts, whereas *L. heterotoma* was more likely to parasitize *D. melanogaster* and *D. obscura* species group. In contrast, PDI for *G. kimorum* did not differ from null expectations. This counterintuitive outcome for *G. kimorum* likely arose because *G. kimorum* was reared exclusively from *in situ* rotting fruit, a substrate in which >91% of available hosts were *D. suzukii*. Since *G. kimorum* does not appear to forage on rotting fruits (Seehausen et al. 2022; Giorgini et al. 2024), where non-*suzukii* hosts were much more abundant in our study, the null model treated its single-host use as the expected outcome for that sampling context. Consequently, the model judged *G. kimorum*’s exclusive association with *D. suzukii* as indistinguishable from frequency-dependent host use. This illustrates an important nuance when interpreting specialization metrics: for specialists confined to particular resource environments, strong habitat filtering can make use of a single host species appear consistent with null predictions (frequency-dependent host species use), even when the true host range of the consumer is extremely narrow (in this case, restricted to a single host species).

It is interesting to note that nearly all of the drosophilids present in the samples are non-native species, with the exception of a minority (11%) of the *D. obscura* group (*D. affinis*), as revealed by DNA barcoding. Most of the *obscura* group samples were made up of the non-native *Drosophila subobscura*. Because only a relatively small subset of emerged host samples were DNA barcoded, it is possible that the *D. obscura* group puparia parasitoids were reared from also included other native species (e.g. *D. athabasca*, *D. pseudoobscura*) in the *D. obscura* species group. The development of specimen-level molecular diagnostic tools for host-parasitoid associations (e.g. Gariepy et al. 2014) could resolve species-level interactions in more detail. Regardless, in this broader context where much of the drosophilid community is non-native (having limited conservation importance; Heimpel et al. 2024) and has no obvious cultural or economic importance to humans, not all of the so-called “non-target” parasitism by *L. japonica* would necessarily be considered a negative environmental consequence of the unintentional introduction of this parasitoid (Rossi Stacconi et al. 2025). *Drosophila melanogaster*, for example, is a non-native nuisance pest of human dwellings in North America (Keller 2007) and the use of this host species by *L. japonica* may cause apparent competition between *D. melanogaster* and *D. suzukii* that helps augment biological control of *D. suzukii*. Frequent parasitism of *D. melanogaster* by *L. japonica* may also provide an alternative host during seasonal or spatial gaps in the availability of *D. suzukii*. However, it is possible that our methodology was particularly likely to sample and rear out non-native species. Sampling of additional substrates (e.g., naturally occurring fungi) that may contain other native species of drosophilids is still required to better understand whether *L. japonica* may have negative effects on native drosophilid species (Rossi Stacconi et al. 2025).

### Host sharing with other parasitoids of drosophilid larvae

Similar to the drosophilid community, the parasitoids reared were almost entirely non-native species. The most common parasitoid attacking drosophilids other than *D. suzukii* was the cosmopolitan species *L. heterotoma* (Quicray et al. 2023), which was most commonly associated with *D. melanogaster* but was also the dominant parasitoid of *D. obscura* group larvae and was the only parasitoid reared from *D. immigrans* larvae. *Asobara* cf. *rufescens*, which is of unknown biogeographic origin but whose arrival in North America pre-dates *L. japonica* and *G. kimorum* (Abram et al. 2020), was reared from all hosts except *D. immigrans*. The fact that *A.* cf. *rufescens* was often reared from *D. suzukii* puparia in *in situ* rotting fruit was initially surprising, given that it is rare to rear this parasitoid species (and other *Asobara* spp. in invaded regions) on this host in the laboratory (Chabert et al. 2012; Abram et al. 2020). However, a laboratory study conducted in parallel to the current study revealed that *A.* cf. *rufescens* is likely acting as a kleptoparasitoid of *D. suzukii* larvae when they are already parasitized by *L. japonica* (Moon et al. 2025).

The two parasitoids that most often share a host in common in this study system appear to be the congeners *L. heterotoma* and *L. japonica* parasitizing *D. melanogaster*. Given that *L. japonica* arrived in the study region between 2012 and 2016 (Gariepy et al. 2024), this is a relatively ‘new’ indirect ecological interaction between parasitoid species sharing a common host that deserves further attention in terms of its consequences for parasitoid-drosophilid food webs and indirect consequences for biological control of *D. suzukii*. A recently-developed multiplex PCR assay that simultaneously screens for *L. heterotoma*, *L. japonica*, and *G. kimorum* (Gariepy et al. 2024) may also be useful in this context – particularly to detect in-host competition (multiparasitism) of *D. melanogaster* and *D. suzukii* by these parasitoids.

Because the methodology in this study was specifically designed to investigate larval parasitoid communities, parasitoid-host relationships of the pupal parasitoids known to be present in the study region were not captured (see Capko et al. 2024). As demonstrated previously, these pupal parasitoids can potentially act as facultative hyperparasitoids or competitors of larval parasitoids attacking *D. suzukii* (and other drosophilids) (Hougardy et al. 2022; Lisi et al. 2025), and thus may further affect trophic interactions in complex ways that should be investigated in future field studies.

## Conclusions

The results of this study strengthen the current consensus that laboratory host range testing as part of classical biological control programs is informative, but conservative. This study focused on non-native larval parasitoids of *D. suzukii*, the more host-specific *G. kimorum* can parasitize *D. melanogaster* in highly artificial laboratory trials (Girod et al. 2018b; Seehausen et al. 2020; Daane et al. 2021), but did not in the field under the conditions investigated here. The less host-specific *L. japonica* attacked both groups of drosophilids predicted to be at most risk in laboratory trials, but attacked one (*D. obscura* species group) at much lower levels than the other (*D. melanogaster* species group) in the field. Thus, although *L. japonica* was previously eliminated from consideration for intentional classical biological control introductions due to its oligophagy, this study suggests that – at least with respect to the community of drosophilids in decaying fruit in south coastal British Columbia – it is now the most important larval parasitoid of the major invasive insect pest *D. suzukii*, and may be unlikely to have significant population-level impacts on native non-target drosophilids under realistic field conditions. Thus, *L. japonica* represents a case of a former candidate classical biological control agent that may have an overall positive benefit-risk balance and perhaps, with the benefit of hindsight, should not have been eliminated from consideration for intentional introductions (*sensu* Hinz et al. 2014). Continuing to study its host range and biological control impact on *D. suzukii* in other regional contexts (e.g., where non-target drosophilid communities are made up of more native species) and habitats, applying some of the same methods developed here, could help the balance of its benefits and risks come into clearer focus and inform whether it should be introduced to other areas of the world or used for augmentative biological control where it is already adventive (Rossi Stacconi et al. 2025).

## Acknowledgements

We thank Clarissa Capko, Victoria Makovetski, and Allison Bruin for technical assistance in the field and laboratory, and to Emily Grove for the artwork used in Figure 5. Thanks to George Heimpel, Tim Haye, Peter Mason, and Chia-Hua Lue for helpful comments on previous versions of the manuscript. Thanks to Blas Lavandero for advice about food web visualization.

## Author contribution statement

PKA conceived research and wrote the manuscript. PKA and JM analyzed data and created the figures. PKA, JF, JM, JT, and TDG conducted field and laboratory work. TDG contributed reagents and analytical tools. TDG and PKA acquired funding. All authors read and approved the manuscript.

