## Supplementary material for "The realized host specificity of *Leptopilina japonica* and *Ganaspis kimorum*, adventive larval parasitoids of the invasive *Drosophila suzukii*": Online Resource 1 (from another version of this preprint: version 1, 2025-12-24)

Paul K. Abram[1*], Jessie Moon[1,2], Jessica L. Fraser[1,3,4], Jason Thiessen[1], Tara D. Gariepy[5]
[1]Agriculture and Agri-Food Canada, Agassiz Research and Development Centre, Agassiz, BC, Canada
[2]Simon Fraser University, Department of Biology, Burnaby, BC, Canada
[3]Université de Montréal, Département de sciences biologiques, Montréal, QC, Canada
[4]Agriculture and Agri-Food Canada, Saint-Jean-sur-Richelieu Research and Development Centre, Saint-Jean-sur-Richelieu, QC, Canada
[5]Agriculture and Agri-Food Canada, London Research and Development Centre, London, ON, Canada

### Methods S1

*Notes on sampling*

Dropped, rotting fruit (cultivated raspberries and blackberries; not treated with insecticides) was only reliably present for limited seasonal periods at experimental farm sites with large abundances of cultivated fruit (one in 2022, two in 2023). At backyard and forested sites, dropped fruit was not ever present in high enough quantities for sampling in either year due to rapid desiccation or fruit predation by other organisms (e.g., slugs) (P. Abram, personal observations). Mushroom bait traps – designed to attract the fungivorous community of drosophilids and their parasitoids – were deployed in 2022, but because they frequently desiccated and no parasitoids emerged, the results are not included in this study.

*Identification of drosophilid puparia*

Because there is not a published identification key to Drosophilidae of the study region, and there were too many specimens to have all of them identified individually by DNA barcoding or taxonomic experts, puparial morphology was used as the primary identification tool

(Abram et al. 2022b) (Figure S1). Based on published resources (Miller 2017; Paiero 2017; deWaard 2017) and previous unpublished work by our group, common drosophilids associated with rotting fruit in the study area were expected to possibly include: (1) *D. suzukii* and the two other non-native species in the *Drosophila melanogaster* species group (*Drosophila melanogaster* Meigen and *D. simulans* Sturtevant); (2) native (e.g., *Drosophila affinis* Sturtevant, *D. athabasca* Sturtevant & Dobzhansky, *D. pseudoobscura* Frolova) and non-native (*Drosophila subobscura* Collin) drosophilids in the *D. obscura* species group; and (3) other non-native species such as *Drosophila immigrans* Sturtevant. Puparia of *D. immigrans* and *D. suzukii* can readily be identified to species and separated from other species and species groups based on morphology (Abram et al. 2022b). Puparia of *D. melanogaster* and *D. simulans* (both in the *melanogaster* species group) cannot be readily differentiated based on puparial morphology. Similarly, species in the *obscura* species group can be distinguished from other species groups but are not distinguishable to the species level. Identifications of a subset of puparia were validated with DNA barcoding.

To validate the puparial identifications and clarify which species may be represented within the species groups (*melanogaster*, *obscura*) delimited by puparia identifications, DNA barcoding was conducted of a subset of emerged adult flies whose puparia were identified as *D. immigrans*, *D. obscura* species group, and *D. melanogaster* species group from the second year of the study (2023). *Drosophila suzukii* puparia were not selected due to their highly distinct puparia morphology and the fact that both male and female adult flies are morphologically distinct from other drosophilid fauna in British Columbia (Abram et al. 2022b). Initially, a total of 399 flies were selected from among samples of adult flies to represent a range of sampling dates from all three 2023 study sites, as well as both sampling methods. Among these, 365 yielded high-quality DNA barcodes that allowed species-level identifications (dropped fruit: 70 samples; sentinel bait traps: 295 samples). Samples came from across the seasonal range of collection dates and from all three sites to maximize the chance of detecting different species. DNA was extracted from individual flies using a non-destructive Chelex extraction, and amplification and bi-directional sequencing of the DNA barcode region was as described by Gariepy et al. (2024). Individuals were identified to species level using the Animal Species Library and the Validated Canadian Arthropod Library in the BOLD ID engine (https://id.boldsystems.org/).

Identifications of puparia based on morphology were highly accurate, as confirmed by DNA barcoding. More than 99% (114/115) of the sampled puparia identified as *D. immigrans* were identified as such by DNA barcoding; however, one was actually *D. suzukii*. Similarly, 98% (147/150) of puparia identified as *D. melanogaster* species group belonged to the non-native species *D. melanogaster* according to DNA barcoding. *Drosophila simulans* (the other member of the *D. melanogaster* group known to be present in BC) was not ever detected among the subsample of flies barcoded. Two puparia identified as *D. melanogaster* were actually *D. suzukii*, likely because the spiracles had broken off making the identification more difficult. One was identified as *Scaptomyza* and may have been contamination. 100% (all 100) of the puparia identified as *D. obscura* species group were correctly identified. Of these, 89% were the non-native species *D. subobscura* and 11% were the native species *D. affinis*.

**Table 3.** Collections of fresh, ripening fruit in 2023 from different species of fruiting plants, showing the sample weight and the number of emerging *Ganaspis kimorum* and *Leptopilina japonica.*

| **Site** | **Date** | **Species of fruiting plant** | **Fruit sample weight (g)** | **# *G. kimorum*** | **# *L. japonica*** |
|---|---|---|---|---|---|
| Experimental Farm #1 | 26-Jun | *Rubus idaeus* | 632.8 | 2 | 12 |
| | 6-Jul | *R. idaeus* | 628.7 | 1 | 3 |
| | 13-Jul | *R. idaeus* | 687.3 | 9 | 3 |
| | 18-Jul | *R. idaeus* | 381.6 | 0 | 5 |
| | 27-Jul | *R. idaeus* | 706.5 | 2 | 35 |
| | 2-Aug | *R. armeniacus* | 695.2 | 1 | 153 |
| | 10-Aug | *R. armeniacus* | 567.6 | 44 | 109 |
| | | *R. armeniacus* | 490.3 | 28 | 208 |
| | 15-Aug | *R. armeniacus* | 258.2 | 10 | 5 |
| | | *R. idaeus* | 236.3 | 22 | 28 |
| Experimental Farm #2 | 26-Jun | *Prunus avium* | 537.5 | 0 | 0 |
| | 5-Jul | *P. avium* | 777.9 | 0 | 8 |
| | 13-Jul | *R. idaeus* | 666.6 | 0 | 3 |
| | 19-Jul | *R. idaeus* | 741.6 | 0 | 1 |
| | 27-Jul | *R. idaeus* | 567 | 66 | 62 |
| | 4-Aug | *R. armeniacus* | 582.6 | 85 | 135 |
| | | *R. idaeus* | 652.5 | 0 | 1 |
| | 9-Aug | *R. armeniacus* | 712.1 | 0 | 0 |
| | | *R. idaeus* | 610.9 | 38 | 34 |
| | 16-Aug | *R. armeniacus* | 822 | 0 | 2 |
| | | *R. idaeus* | 538.8 | 2 | 9 |
| | 26-Jun | *Prunus avium* | 230.5 | 0 | 0 |
| Mid-elevation forest | 26-Jun | *Sambucus racemosa* | 410.9 | 0 | 0 |
| | 5-Jul | *S. racemosa* | 274.3 | 0 | 1 |
| | 13-Jul | *S. racemosa* | 234.7 | 0 | 0 |
| | 19-Jul | *S. racemosa* | 252.6 | 2 | 0 |
| | 27-Jul | *R. armeniacus* | 622.8 | 9 | 0 |
| | 4-Aug | *R. armeniacus* | 735 | 4 | 0 |
| | 9-Aug | *R. armeniacus* | 563.8 | 0 | 14 |
| | 16-Aug | *R. armeniacus* | 339.5 | 0 | 12 |

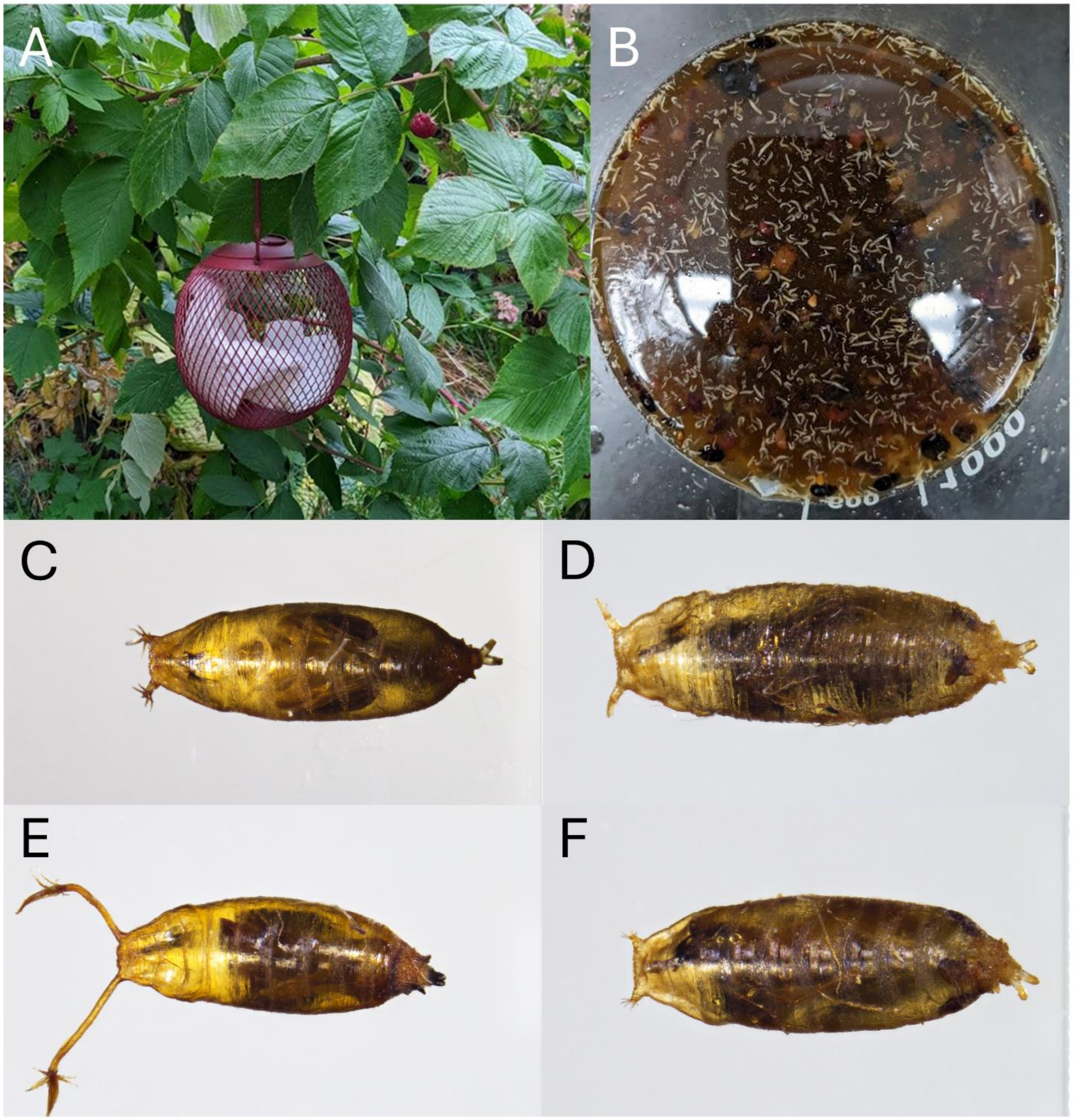


**Figure 6.** Bait trapping and drosophilid puparium extraction. (A) A metal "NO-NO" bird feeder trap containing fruit bait; (B) Brown sugar brine with drosophilid larvae from rotting fruit floating to the top; (C) Example of a *Drosophila suzukii* puparium; (D) Example of a *D. melanogaster* species group (other than *D. suzukii*) puparium; (E) Example of a *D. immigrans* puparium; (F) Example of a *D. obscura* species group puparium.

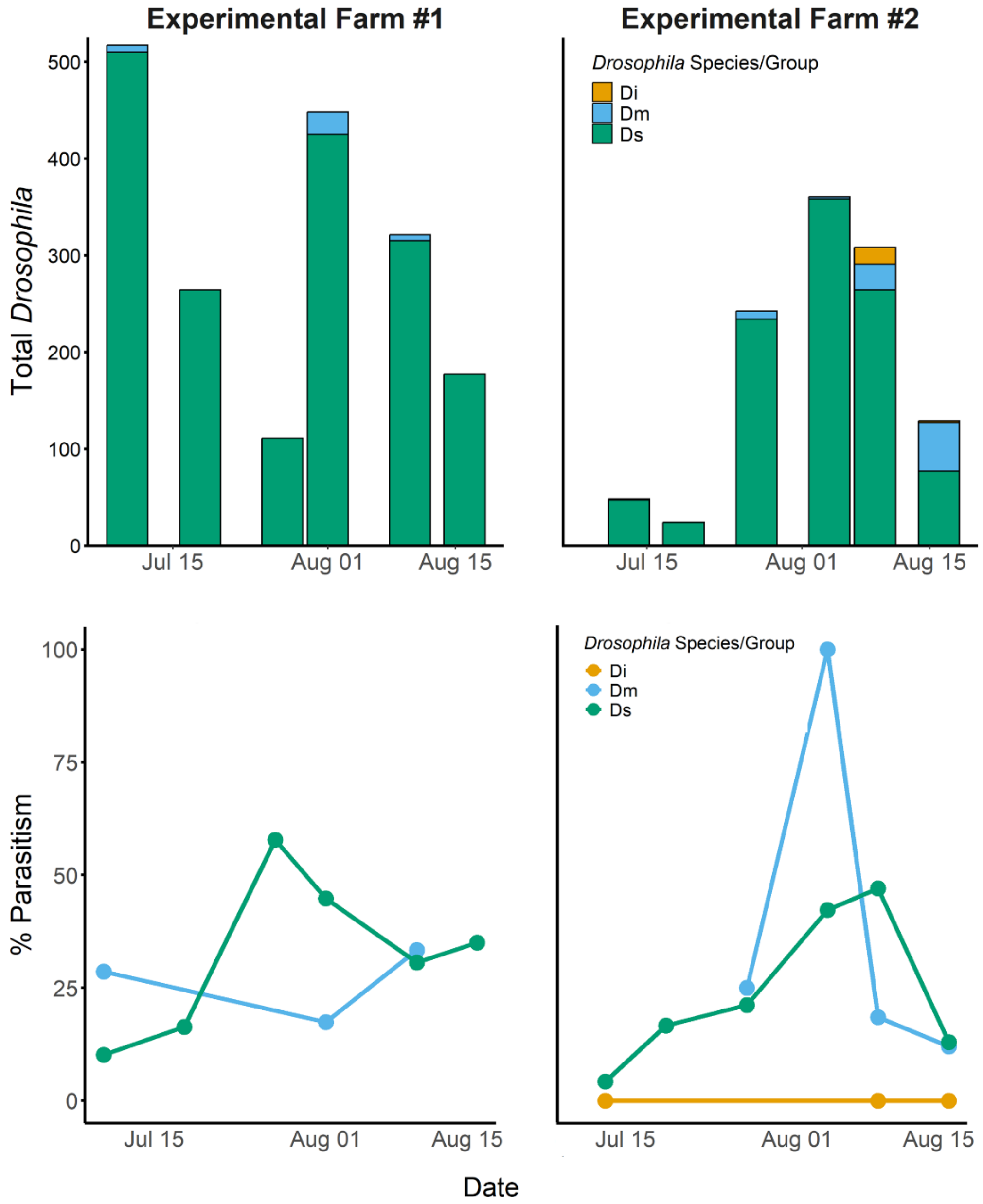


**Figure 7.** The total number of *Drosophila* puparia collected from rotting fruits in the second year of the study (2023) on each collection date at each site (upper panels) and the percentage of the puparia of each species or species group from which parasitoids emerged (lower panels). Species abbreviations: Di – *Drosophila immigrans*; Dm – *D. melanogaster* species group other than *D. suzukii*; Ds – *Drosophila suzukii,*

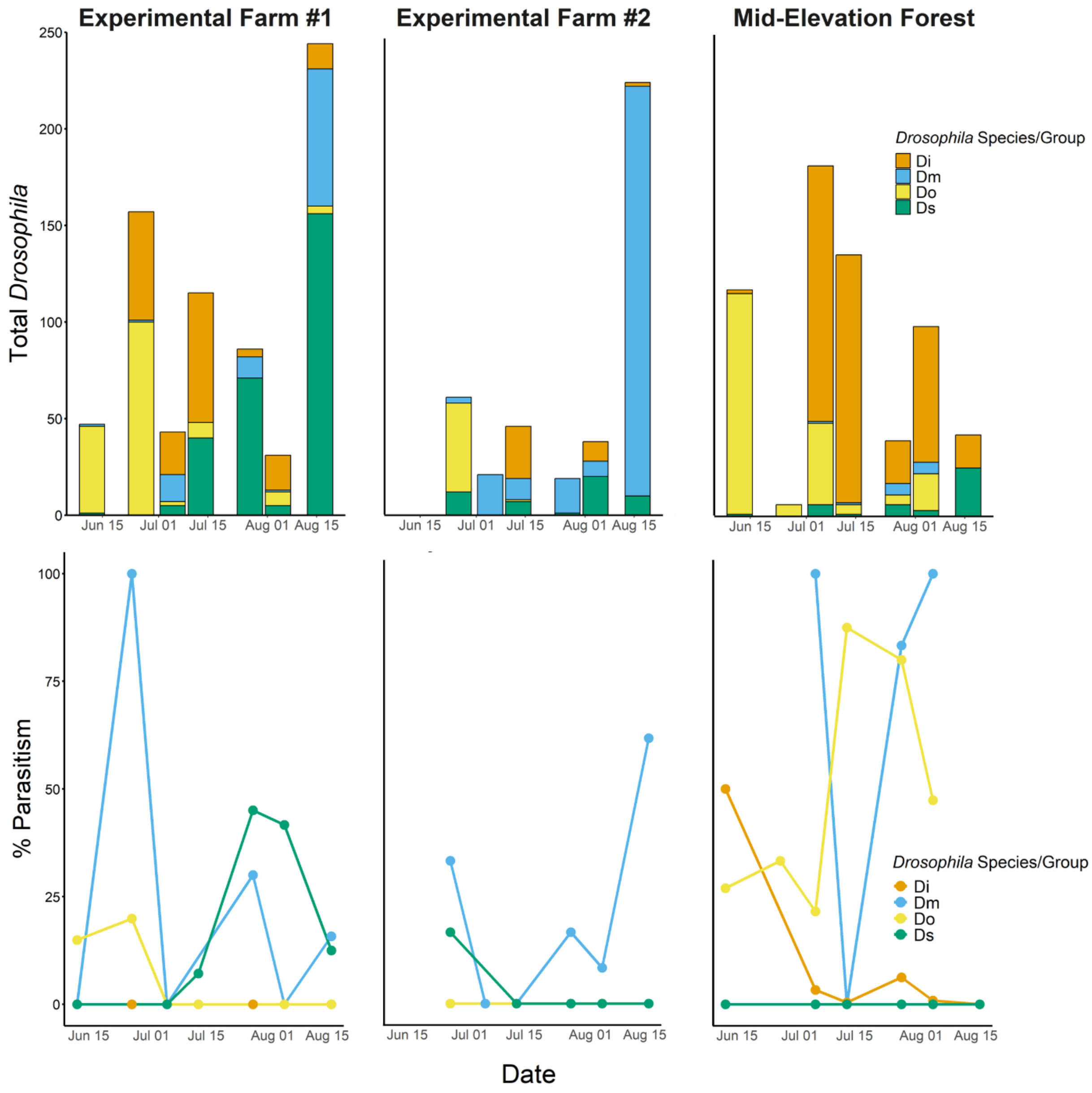


**Figure 8.** The total number of *Drosophila* puparia collected from sentinel fruit bait traps in the second year of the study (2023) on each collection date at each site (upper panels) and the percentage of the puparia of each species or species group from which parasitoids emerged (lower panels). Species abbreviations: Di – *Drosophila immigrans*; Dm – *D. melanogaster* species group other than *D. suzukii*; Do – *D. obscura* species group; Ds – *Drosophila suzukii,*

**References S1**

Abram PK, Wang X, Hueppelsheuser T, Franklin MT, Daane KM, Lee JC, et al (2022b) A coordinated sampling and identification methodology for larval parasitoids of spotted-wing drosophila. J Econ Entomol 115:922–942.

deWaard J (2017) Centre for Biodiversity Genomics (BIOUG). Version 9.6. Univ Guelph. Occurrence dataset. https://doi.org/10.5886/qzxxd2pa

Gariepy TD, Abram PK, Adams C, Beal D, Beers E, Beetle J, et al (2024) Widespread establishment of adventive populations of *Leptopilina japonica* in North America and development of a multiplex PCR assay to identify key parasitoids of *Drosophila suzukii*. NeoBiota 93:63–90.

Miller ME, Marshall SA, Grimaldi DA (2017) A review of the species of *Drosophila* and genera of Drosophilidae of northeastern North America. Can J Arthropod Identif 31.

Paiero S (2017) Univ Guelph Insect Collection (DEBU). Version 2.2. Univ Guelph. Occurrence dataset. https://doi.org/10.5886/4pp2vt5a
